# Activin A mediated KAT8 expression induces ferroptosis during *Mycobacterium tuberculosis* infection

**DOI:** 10.1101/2025.03.06.641808

**Authors:** Bijewar Ashish Satish, Smriti Sundar, Raju S Rajmani, Kithiganahalli Narayanaswamy Balaji

**Affiliations:** Department of Microbiology and Cell Biology, Indian Institute of Science, Bengaluru, Karnataka, India; Centre for Infectious Disease Research, Indian Institute of Science, Bengaluru, Karnataka, India

**Author notes:** To whom correspondence should be addressed: Dr. Kithiganahalli Narayanaswamy Balaji, Department of Microbiology and Cell Biology, Indian Institute of Science, Bangalore 560012, Karnataka, India.

**Keywords:** Activin A, *Mycobacterium tuberculosis*, Ferroptosis, KAT8, NRF2, HO-1

## Abstract

Activin A, a secretory glycoprotein, has been shown to be upregulated in TB patients, and its levels are correlated with disease severity. Here, we identify a functional role for activin A and the downstream SMAD2/3 signalling in Mtb pathogenesis and dissemination. Molecular assays, including ChIP and loss-of-function analysis, demonstrated that Activin A regulates the expression of lysine acetyltransferase (KAT) 8, which in turn regulates HO-1 levels and Mtb-induced ferroptosis. Mechanistically, we identify KAT8-mediated acetylation of NRF2 during Mtb infection, leading to enhanced nuclear availability of the transcription factor and increased expression of HO-1. Finally, utilizing an *in vivo* mouse model of TB, we show that the pharmacological inhibition of activin A receptor and KAT8 restricts Mtb burden, dissemination and ameliorates TB pathology. Thus, we report a novel role for activin A in regulating NRF2 localisation and outline its potential consequences during TB.

## Introduction

Tuberculosis (TB), caused by the intracellular pathogen *Mycobacterium tuberculosis* (Mtb), reprogrammes distinct cellular events to evade host immunity, thereby enhancing pathogen dissemination and survival (1, 2). Cellular reprogramming of macrophages, a primary site for Mtb infection, dictates survival within the host environment by resulting in impaired macrophage effector functions, including the active blockade of phagosome lysosome fusion, inhibition of apoptosis, and inhibition of antigen presentation (1, 3). Studies have identified the role of differentially regulated proteins, including immunomodulators and immunoregulatory transcription factors, in regulating macrophage effector functions during Mtb infection (4–6). Additionally, the elevated levels of distinct cytokines, such as IL-10, Type I-IFN, and IL-13, found in the serum of TB patients have also been reported to regulate macrophage function and contribute to disease severity (6, 7).

To our interest, a recent study identified an enhanced expression of activin A in the serum of TB patients and indicated their role in disease exacerbation (8). Activins are multi-functional cytokines widely implicated in performing pleiotropic roles in physiological and pathological processes such as cell proliferation, inflammation, and tissue fibrosis (9–11). Activin A, a homodimer of inhibin βA subunits, binds to surface type II activin receptors (ACTRII) and induces the recruitment and phosphorylation of distinct type I activin receptors (ACTRI) such as activin receptor-like kinases (ALK) 4, ALK2 and ALK7. The phosphorylation of ACTRI leads to an activation of its kinase activity. Once activated, ACTRI phosphorylates Mothers against decapentaplegic homolog (SMAD) 2 and 3, forming a heteromeric complex with SMAD 4. The SMAD complex then translocates to the nucleus and regulates downstream gene expression (9, 10). Activin A has been identified to play crucial roles in regulating immune cell functions during infections and autoimmune disorders. For instance, activin A levels correlate with increased pathogen burden, pro-inflammatory cytokine secretion, and oxidative stress (12–14). However, the precise role of activin A and the functional relevance of its downstream signalling events during intracellular infections, including TB, remains largely unknown.

Cell death modalities, resulting from complex host-pathogen dialogues, have been widely implicated in regulating Mtb pathogenesis. Particularly, necrotic lesions observed during TB contribute to the spread of Mtb, attributed to the extracellular growth of bacilli released in tissues (15–17). While specific factors regulating the process and the interplay of different modes of cell death during TB remain incompletely understood, emerging evidence implicates a key role of ferroptosis in contributing to disease pathology. Ferroptosis, an iron-dependent programmed-necrotic cell death, is driven by an accumulation of oxidative species and lipid peroxides, leading to membrane damage and concomitant rupture (18). Mtb infection has been reported to increase the labile iron pool (LIP), impair antioxidant machinery and reduce levels of Glutathione peroxidase (GPX) 4 leading to ferroptosis in macrophages (19–21). In this context, delineation of the molecular mechanisms regulating ferroptosis during TB will assist in developing therapeutic strategies targeting this key node of Mtb pathogenesis.

To our interest, ALK4-mediated signalling was reported to regulate ferroptosis in renal epithelial cells by modulating the expression of the master gene regulator, Nuclear factor erythroid 2-related factor (NRF) 2 (22). Upon distinct stimuli, including post-translational modifications, NRF2 is translocated to the nucleus, where it binds at the antioxidant-responsive elements (ARE) present on the promoter of its target genes and regulates their expression. In this context, acetyltransferases, among others, have been identified to play crucial roles in regulating NRF2 function by modulating its nucleocytoplasmic localisation (23). Notably, lysine acetyltransferase (KAT) 8 can regulate transcriptional activity of NRF2 by acetylating it and enhancing its nuclear availability (24). While events regulating NRF2 localisation during Mtb infection remain largely unknown, NRF2 expression and its downstream effectors have been widely implicated during TB. Specifically, the levels of NRF2-mediated expression of the heme catabolising enzyme, heme oxygenase (HO)-1, have been identified to execute Mtb pathogenesis. HO-1 expression has also been correlated with disease severity in TB, with reduced levels observed upon successful TB treatment and sustained levels observed in those who experienced treatment failure or relapse (25). To our interest, HO-1 inhibition has also been reported to control mycobacterial infection in mice models of TB infection, which was rescued upon exogenous iron supplementation, indicating a crucial role for HO-1 mediated heme catabolism in mycobacterial pathogenesis (26, 27). While HO-1 has been widely implicated in inducing ferroptosis in distinct models of study, its impact on regulating ferroptosis during Mtb infection remains enigmatic. Thus, we were encouraged to explore the effect of activin A signalling on NRF2-mediated regulation of HO-1 levels during mycobacterial infection.

We identify that Mtb induces the robust activation of activin A-mediated SMAD2/3 signaling. Concurrently, we establish a role for activin-induced signaling in regulating ferroptosis during Mtb infection by regulating the nuclear retention of NRF2 and therein levels of HO-1. Importantly, we show that the pharmacological inhibition of this process leads to reduced Mtb survival and reduced dissemination in mice models of TB infection.

## Results

### Activin A regulates SMAD2/3 activation during Mtb infection

Cellular reprogramming events due to altered immune signalling play crucial roles in the outcome of Mtb infection(28–30). To determine the role of activin A in regulating mycobacterial pathogenesis, we assessed the levels of activin ligands, receptors, and the downstream effectors in both *in vitro* and *in vivo* scenarios. We observed that Mtb infection resulted in the robust upregulation of activin A (*Inhba)* and activin B *(Inhbb)* at transcript levels (Figure 1A). Enhanced levels of phosphorylated SMAD 2/3 (Figure 1B) and expression of ALK4 (Figure 1C) were also observed in cells infected with Mtb. These results were corroborated in an *in vivo* model of TB (Figure 1D) wherein elevated levels of activin ligands, phosphorylated SMAD 2/3 and ALK4 were observed in RNA and protein extracted from the lungs of infected mice (Figure 1E-G). Further, the addition of follistatin (FST), a glycoprotein that binds to activin A and blocks its biological activity, compromised Mtb-induced enhanced phosphorylation of SMAD 2/3 validating a key role for the ligand in mediating the downstream effectors of the signalling (Figure 1H). The addition of the ALK4 inhibitor, SB-431542, that inhibits cellular SMAD2 phosphorylation also reduced the levels of phosphorylated SMAD 2/3, indicating a key role for the cognate receptor in the activation of activin signalling during Mtb infection (Figure S1A). Similarly, reduced levels of phosphorylated SMAD 2/3 were also validated in infected mice administered with SB-431542 when assessed in lung homogenates (Figure S1B).

**Figure 1:**
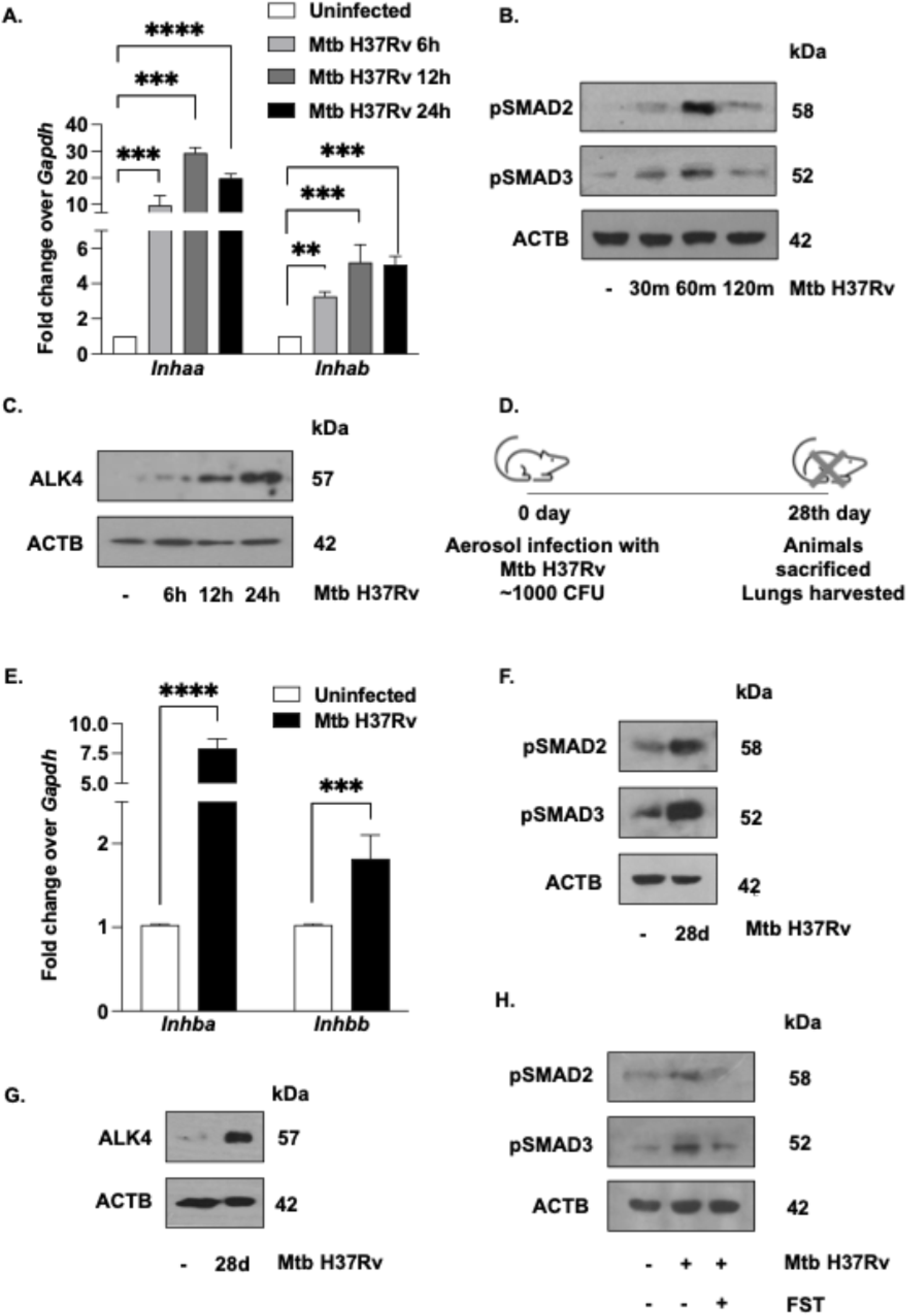
Mtb infection elevates Activin Signalling *in vitro* and *in vivo*. **(A)** Mouse macrophages were infected with Mtb H37Rv for indicated time points and assessed for transcript levels of activin ligands *Inhba* (Activin A), *Inhbb* (Activin B) by qRT-PCR. **(B, C)** Mouse macrophages were infected with Mtb H37Rv for indicated time points and assessed for **(B)** phosphorylation of SMAD2/3 **(C)** protein levels of ALK4 by immunoblotting. **(D)** Schematic representing the time course of a mouse model of Mtb infection **(E-G)** BALB/c mice were aerosol-infected (∼1000 colony-forming units [CFU]) with Mtb H37Rv for 28 d and **(E)** transcript levels of activin ligands were assessed in whole lungs by qRT-PCR **(F)** phosphorylation of SMAD2/3 were assessed in lung homogenate of uninfected and infected mice using immunoblotting **(G)** protein expression of ALK4 were assessed in lung homogenate of uninfected and infected mice using immunoblotting. **(H)** Mouse macrophages were pre-treated with Follistatin (FST) and infected with Mtb H37Rv for 60 mins and assessed for phosphorylation of SMAD2/3 by immunoblotting. All qRT-PCR data represents mean ± S.E.M., and immunoblotting data is representative of three independent experiments. *** p < 0.001; ****, p < 0.0001 (Student’s t-test in A and E; GraphPad Prism 10.0). ACTB was utilized as a loading control

### Activin signalling induces ferroptosis during Mtb infection

As Mtb infection induces a robust activation of activin-mediated SMAD2/3 signalling, evaluating its role during infection was imperative. A recent study demonstrated that the perturbation of ALK4/5 receptors in renal epithelial cells inhibits erastin-induced ferroptosis (22). Ferroptosis in macrophages, driven by the accumulation of labile iron pool (LIP) and lipid peroxides, has also been implicated during Mtb infection (19, 21). Thus, we hypothesised a possible correlation between activin signalling and ferroptosis in macrophages during Mtb infection and aimed to investigate its role. To assess the extent of lipid peroxidation, cells were stained with BODIPY 493/503, a dye that stains neutral lipids, and BODIPY 665/676, a dye that stains unoxidized lipids. The ratio between unoxidized and neutral lipids was calculated and represented as an indicator of lipid peroxidation. We observed that the addition of ALK4 inhibitor or depletion of ALK4 in macrophages reduced the accumulation of peroxidized lipids during Mtb infection (Figure 2A, B; S2A, B). Similarly, we observed that the addition of ALK4 inhibitor significantly reduced cellular LIP upon Mtb infection, indicating its role in iron metabolism and, therefore, ferroptosis (Figure 2C). Concomitantly, inhibition of ALK4 also reduced necrotic cell death in Mtb-infected macrophages quantified by the release of lactate dehydrogenase (LDH) into the culture supernatants (Figure 2D). Ferroptosis in macrophages facilitates Mtb release into the extracellular milieu and contributes to mycobacterial dissemination during infection (19). Consistent with this, increased cell death was accompanied by an increase in extracellular bacteria which was subsequently compromised upon inhibition of activin signalling (Figure 2E; S2C), indicating a role of activin signalling in regulating ferroptosis during Mtb infection.

**Figure 2:**
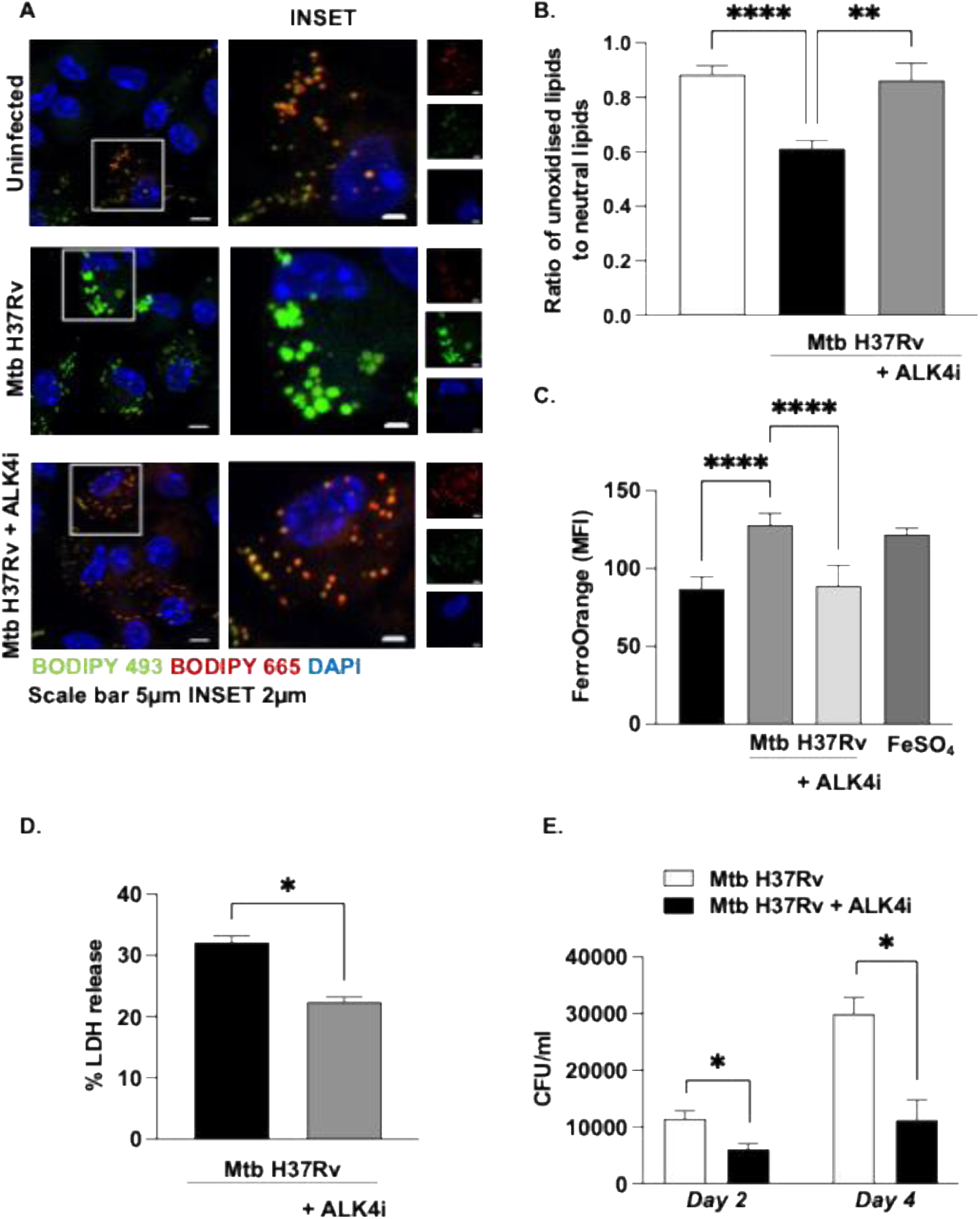
Activin signalling induces ferroptosis during Mtb infection. **(A, B)** Mouse macrophages were pre-treated with ALK4 inhibitor (SB-431542) followed by 24 h infection with Mtb H37Rv. Lipids were stained with BODIPY 493/503 and BODIPY 665/676 to quantify lipid peroxidation and observed by confocal microscopy **(A)** representative images and **(B)** the respective quantification. **(C)** Mouse macrophages were pre-treated with ALK4 inhibitor (SB-431542) followed by 24 h infection with Mtb H37Rv. The labile iron pool was quantified via staining with FerroOrange. **(D)** Necrotic cell death was measured by LDH release assay in mouse macrophages treated with ALK4 inhibitor (SB-431542), followed by Mtb H37Rv infection for 96 h. **(E)** The release of live mycobacteria from necrotic cells was examined by CFU quantification in mouse macrophage pre-treated with ALK4 inhibitor (SB-431542) by plating culture supernatants 48 h and 96 h post-infection. All confocal microscopy data are representative of three independent experiments. *, p< 0.05; **, p< 0.005;; ****, p < 0.0001 (Student’s t-test in D and E; One-way ANOVA in B, C; GraphPad Prism 10.0). SB431542: ALK4 inhibitor. Abbreviations: CTCF, corrected total cell fluorescence; DAPI, 4′,6-diamidino-2-phenylindole; CFU, colony forming unit.

### Activin signalling regulates the nuclear retention of NRF2 during Mtb infection

Interestingly, activin A and ALK4 have been previously reported to regulate ferroptosis by activating the transcription factor NRF2 (22, 31). NRF2 and its downstream effector HO-1, a rate-limiting enzyme in heme degradation, have been reported to induce iron-dependent ferroptosis by releasing large amounts of free iron within the cell (32, 33). In line with this, we observed that the inhibition of NRF2 significantly reduced the LIP and necrotic cell death of Mtb-infected macrophages (Figure S3A, B). Additionally, the depletion of HO-1 significantly compromised the mycobacterial burden in the culture supernatants of infected macrophages (Figure S3C). Thus, we aimed to assess a potential role for HO-1 in activin signalling-induced ferroptosis during Mtb infection. Our initial studies confirmed the enhanced expression of HO-1 upon Mtb infection (Figure 3A; S3D, E). In concert with our hypothesis, the addition of ALK4 inhibitor significantly compromised Mtb-induced expression of HO-1 in both *in vitro* and *in vivo* models of Mtb infection (Figure 3B). Based on this result, we surmised that activin signalling might regulate NRF2-mediated expression of HO-1 during Mtb infection. Chromatin immunoprecipitation (ChIP) assay confirmed enhanced recruitment of NRF2 on the promoters of HO-1 upon Mtb infection and a concomitant decrease upon the inhibition of ALK4 (Figure 3C). To determine the mechanistic details of altered NRF2 activity, we assessed the levels of NRF2 post-treatment with ALK4 inhibitor during Mtb infection. There was no appreciable alteration in the levels of NRF2 (Figure S3F), indicating that the elevated levels of activin signalling could result in the altered localisation or activity of NRF2 leading to enhanced HO-1 levels. The activation of NRF2 requires its dissociation from its partner, Kelch-like ECH-associated protein (KEAP) 1, followed by its nuclear translocation (34). In this regard, we evaluated the nucleocytoplasmic localisation of NRF2 and observed that Mtb infection results in an increased accumulation of NRF2 within the nucleus of macrophages (Figure S3G, H). Thus, we aimed to assess whether activin signalling regulates NRF2-mediated transcription during Mtb infection by modulating its nuclear localisation and observed a compromised nuclear localisation of NRF2 upon the addition of an ALK4 inhibitor during Mtb infection (Figure 3D, E). These results indicate the possible role for activin signalling in favouring mycobacterial pathogenesis by regulating the nuclear localization of NRF2 and inducing HO-1 dependent ferroptosis.

**Figure 3:**
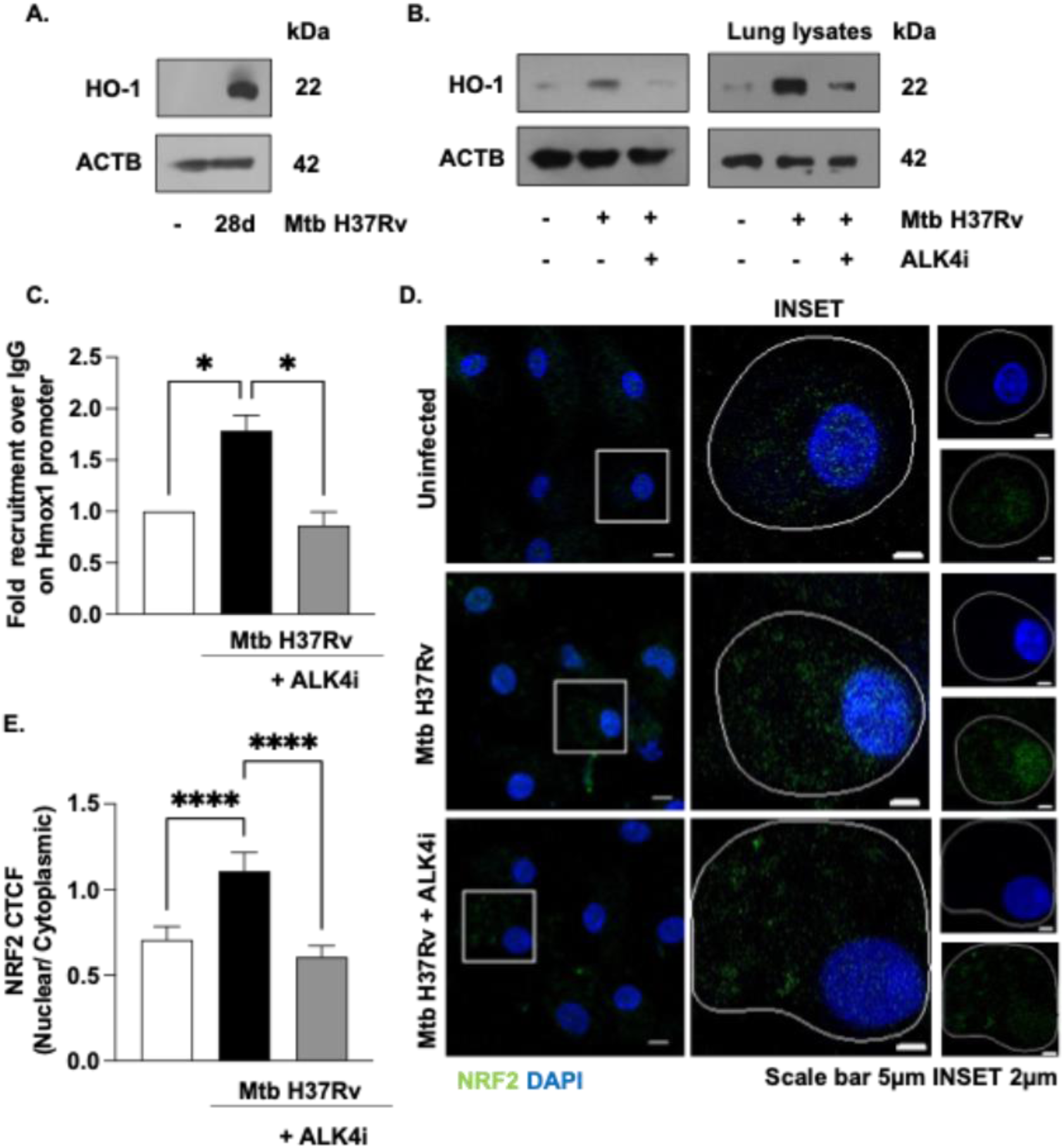
Activin signalling contributes to HO-1 expression during Mtb infection. **(A)** BALB/c mice were aerosol-infected (∼1000 colony-forming units [CFU]) with Mtb H37Rv for 28 d, and protein expression of HO-1 was assessed in the lung homogenate of uninfected and infected mice using immunoblotting. **(B)** Mouse macrophages treated with ALK4 inhibitor (SB-431542) for 1 h followed by 24 h infection with Mtb H37Rv *(left)* and lung lysates from BALB/c mice aerosol-infected (∼1000 colony-forming units [CFU]) with Mtb H37Rv and administered with ALK4 inhibitor (SB-431542) from 15 d p.i *(right)* were assessed for the expression of HO-1 by immunoblotting. **(C)** Mouse macrophages were infected with Mtb H37Rv for 24 h and assessed for the recruitment of NRF2 over the *Hmox1* promoter by ChIP assay. **(D, E)** Mouse macrophages were pre-treated with ALK4 inhibitor (SB-431542) followed by 24 h infection with Mtb H37Rv and assessed for nuclear levels of NRF2 by confocal microscopy **(D)** representative image **(E)** its quantification. Immunoblotting and Immunofluorescence data are representative of three independent experiments. *, p < 0.05; ****, p < 0.0001 (One-way ANOVA in C and E; GraphPad Prism 10.0). ACTB was used as a loading control. SB431542: ALK4 inhibitor. Abbreviations: CTCF, corrected total cell fluorescence; DAPI, 4′,6-diamidino-2-phenylindole.

### KAT8 regulates nuclear retention of NRF2 during Mtb infection

The association of the KEAP1/NRF2 complex is regulated by stimuli such as electrophilic stress or increased levels of reactive oxygen species (ROS) (34). However, it is not clear what molecular cues regulate this process during Mtb infection. Interestingly, post-translational modifications of NRF2 have been increasingly reported in regulating its nucleocytoplasmic localisation (23, 35). Specifically, previous studies have determined that an increased level of NRF2 acetylation enhances its nuclear localization and NRF2-mediated gene expression (23, 36). A recent study implicated the role of KAT8, an acetyltransferase, in the acetylation of NRF2 in non-small cell lung cancer (24). Besides, KAT8 was an intriguing candidate as it has also been implicated in regulating other immunological processes, including apoptosis, autophagy, and cellular ROS levels (37, 38). We found that KAT8 is expressed in *in vitro* macrophages and in the lungs of mice infected with Mtb H37Rv (Figure 4A-C). Immunoprecipitation of NRF2 indicated an enhanced interaction between KAT8 and NRF2 during Mtb infection that was compromised upon the inhibition of ALK4 (Figure 4D). Congruently, we observed enhanced levels of acetylated NRF2 upon Mtb infection that was reduced upon treatment with KAT8 or ALK4 inhibitors (Figure 4E). In line with this, the nuclear localization of NRF2 in Mtb-infected cells was also compromised upon *Kat8* knockdown (Figure 4F, S4A).

**Figure 4:**
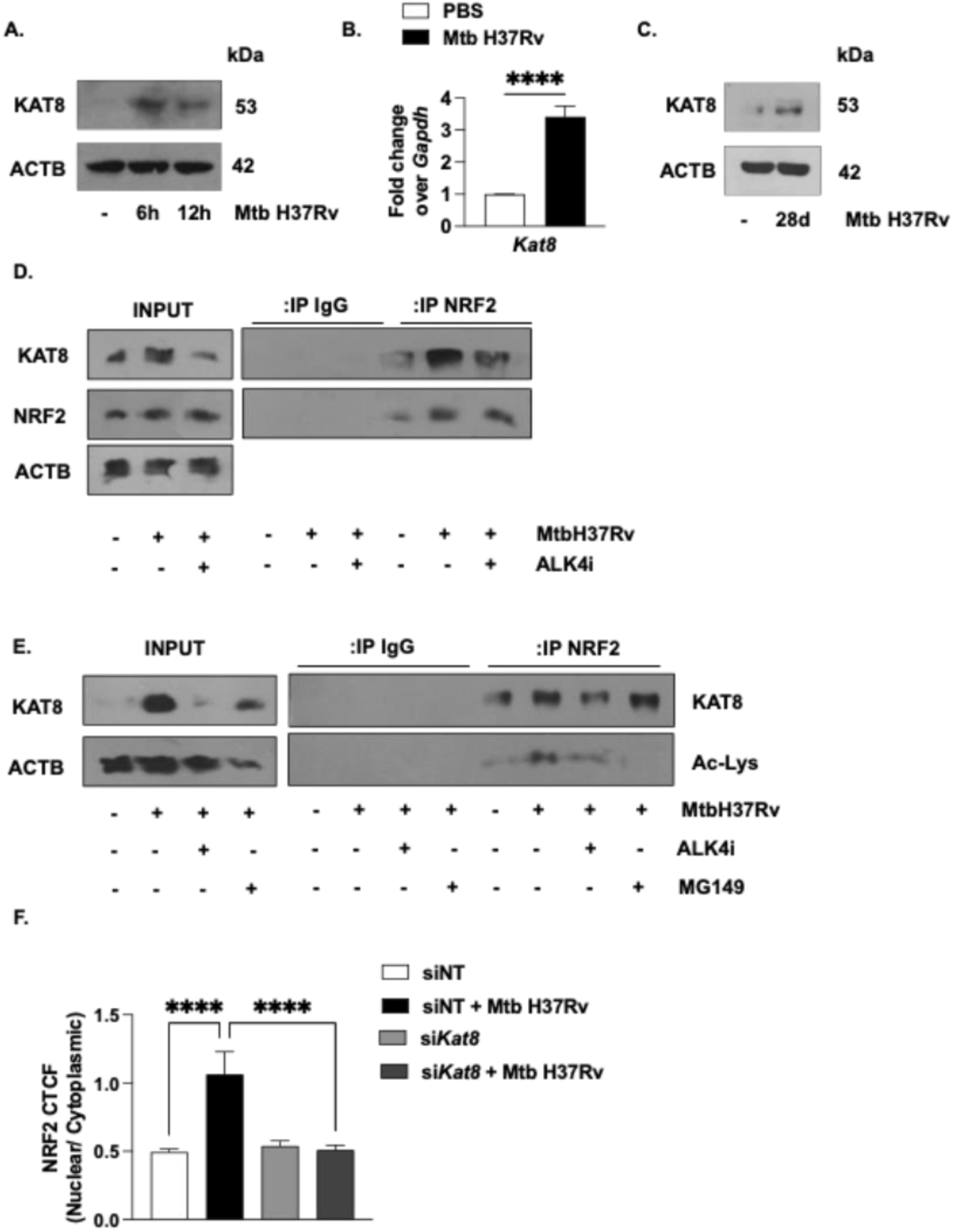
Activin contributes to NRF2 acetylation through KAT8 during Mtb infection. **(A)** Mouse macrophages were infected with Mtb H37Rv for indicated time points and assessed for protein levels of KAT8 using immunoblotting. **(B, C)** BALB/c mice were aerosol-infected (∼1000 colony-forming units [CFU]) with Mtb H37Rv for 28 d, **(B)** transcript levels of *Kat8* were assessed in whole lungs by qRT-PCR **(C)** levels of KAT8 were assessed in lung homogenate of uninfected and infected mice using immunoblotting. **(D)** Mouse macrophages were pre-treated with ALK4 inhibitor (SB431542), infected with Mtb H37Rv for 24 h and assessed for the interaction of KAT8 and NRF2 by immunoprecipitation assay **(E)** Mouse macrophages were pre-treated with ALK4 inhibitor (SB431542) or MG149 (KAT8 inhibitor), infected with Mtb H37Rv for 24 h and assessed for the levels of acetylated lysine on NRF2 and interaction of KAT8 and NRF2 by immunoprecipitation assay. **(F)** Mouse macrophages were transfected with non-targeting or *Kat8* small interfering RNAs. The transfected cells were infected with Mtb H37Rv for 24 h and assessed for nuclear levels of NRF2 by confocal microscopy. All immunoblotting data is representative of three independent experiments. ****, p < 0.0001 (Student’s t-test in B, One-way ANOVA in F; GraphPad Prism 10.0). ACTB was used as a loading control. SB431542: ALK4 inhibitor; MG149: KAT8 inhibitor. Abbreviations: IP, Immunoprecipitation; CTCF, corrected total cell fluorescence.

### Activin-induced KAT8 expression regulates ferroptosis during Mtb infection

Having observed enhanced levels of acetylated NRF2 and its interaction with KAT8 during Mtb infection, we wanted to delineate the mechanism adopted by the activin signalling pathway to regulate this process. Initial experiments indicated that the exogenous addition of activin A to macrophages enhanced the transcript and protein levels of KAT8, suggesting a role for activin signalling in regulating its levels (Figure 5A, B). Further, activin signalling was identified to regulate the expression of KAT8 as the inhibition of ALK4 reduced its levels during Mtb infection (Figure 5C). We then employed a bioinformatics-based approach to scan the promoter of *Kat8*, which indicated the putative binding sites of SMAD3 and 4 (Figure 5D). Subsequently, ChIP assay revealed enhanced recruitment of SMAD3 onto the promoter of *Kat8* during Mtb infection (Figure 5E). Using loss of function analysis using pharmacological inhibition and siRNA mediated silencing, we observed that the depletion of *Kat8* significantly compromised the accumulation of peroxidized lipids during Mtb infection (Figure 5F, S4B) and decreased Mtb released into the extracellular milieu (Figure 5G, S4C). With these observations, we surmised the role of activin signalling and KAT8 in regulating Mtb-induced ferroptosis.

**Figure 5:**
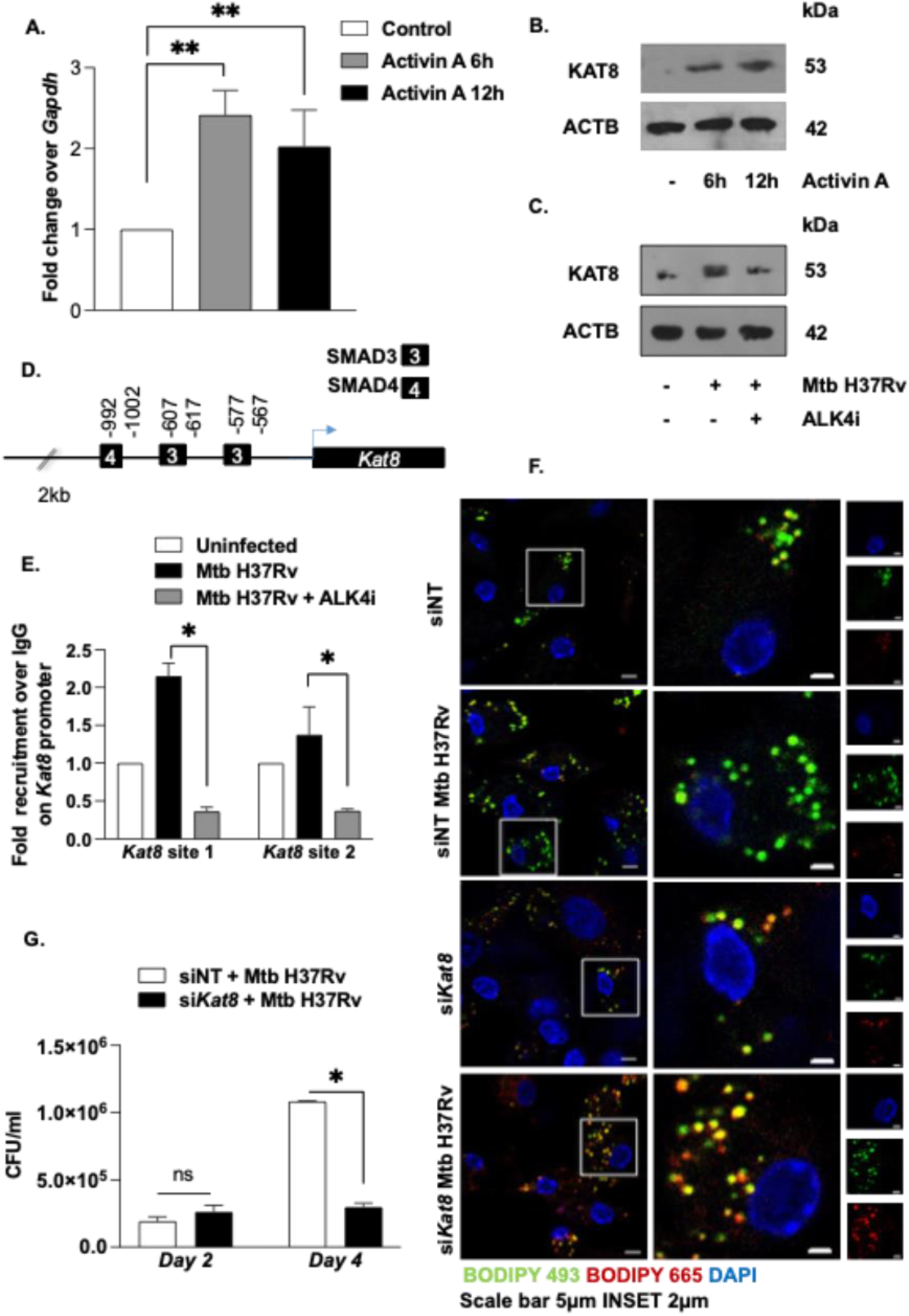
Activin regulates KAT8 levels during Mtb infection. **(A)** Mouse macrophages were treated with activin A for indicated time points and assessed for transcript levels of *Kat8* using qRT-PCR. **(B)** Mouse macrophages were treated with activin A for indicated time points and assessed for protein levels of KAT8 using immunoblotting. **(C)** Mouse peritoneal macrophages were pre-treated with ALK4 inhibitor (SB431542), infected with Mtb H37Rv for 24 h, assessed for protein levels of KAT8 using immunoblotting. **(D)** Bioinformatic assessment of the 2kb upstream sequence of *Kat8* revealed two putative binding sites of SMAD3 **(E)** Mouse macrophages were infected with Mtb H37Rv for 24 h and assessed for the recruitment of SMAD3 over the *Kat8* promoter by ChIP assay. **(F)** Mouse macrophages were transfected with non-targeting or *Kat8* small interfering RNAs. Lipids were stained with BODIPY 493/503 and BODIPY 665/676 to quantify lipid peroxidation and observed by confocal microscopy **(G)** Mouse macrophages were transfected with non-targeting or *Kat8* small interfering RNAs. The release of live mycobacteria from necrotic cells was examined by CFU quantification in transfected cells by plating culture supernatants 48 h and 96 h post-infection. All immunoblotting and confocal microscopy data is representative of three independent experiments. *, p < 0.05; **, p < 0.005 (Student’s t-test in A, E, G; GraphPad Prism 10.0). ACTB was used as a loading control. SB431542: ALK4 inhibitor. Abbreviations: CTCF, corrected total cell fluorescence. NT, non-targeting.

### Activin and KAT8 inhibition ameliorates TB pathology in mice by regulating ferroptosis

Having established the role of activin signalling and KAT8 expression in mediating ferroptosis during Mtb infection, we employed an *in vivo* mouse model of TB to evaluate the effect of their pharmacological inhibition on mycobacterial burden and disease pathology. BALB/c mice were aerosol infected with Mtb and administered ALK4 inhibitor (SB431542) or KAT8 inhibitor (MG149) (Figure 6A). We observed a significant reduction in Mtb burden in the lungs of infected mice upon treatment with the mentioned inhibitors (Figure 6B). Further, we also observed an appreciable decrease in mycobacterial burden in the spleen of infected mice, indicating a role for activin signalling and KAT8 in Mtb dissemination (Figure 6C). The formation of TB-like necrotic lesion in the lungs of infected mice were also significantly compromised upon inhibitor treatment as indicated by the H and E-stained lung sections (Figure 6D).

**Figure 6:**
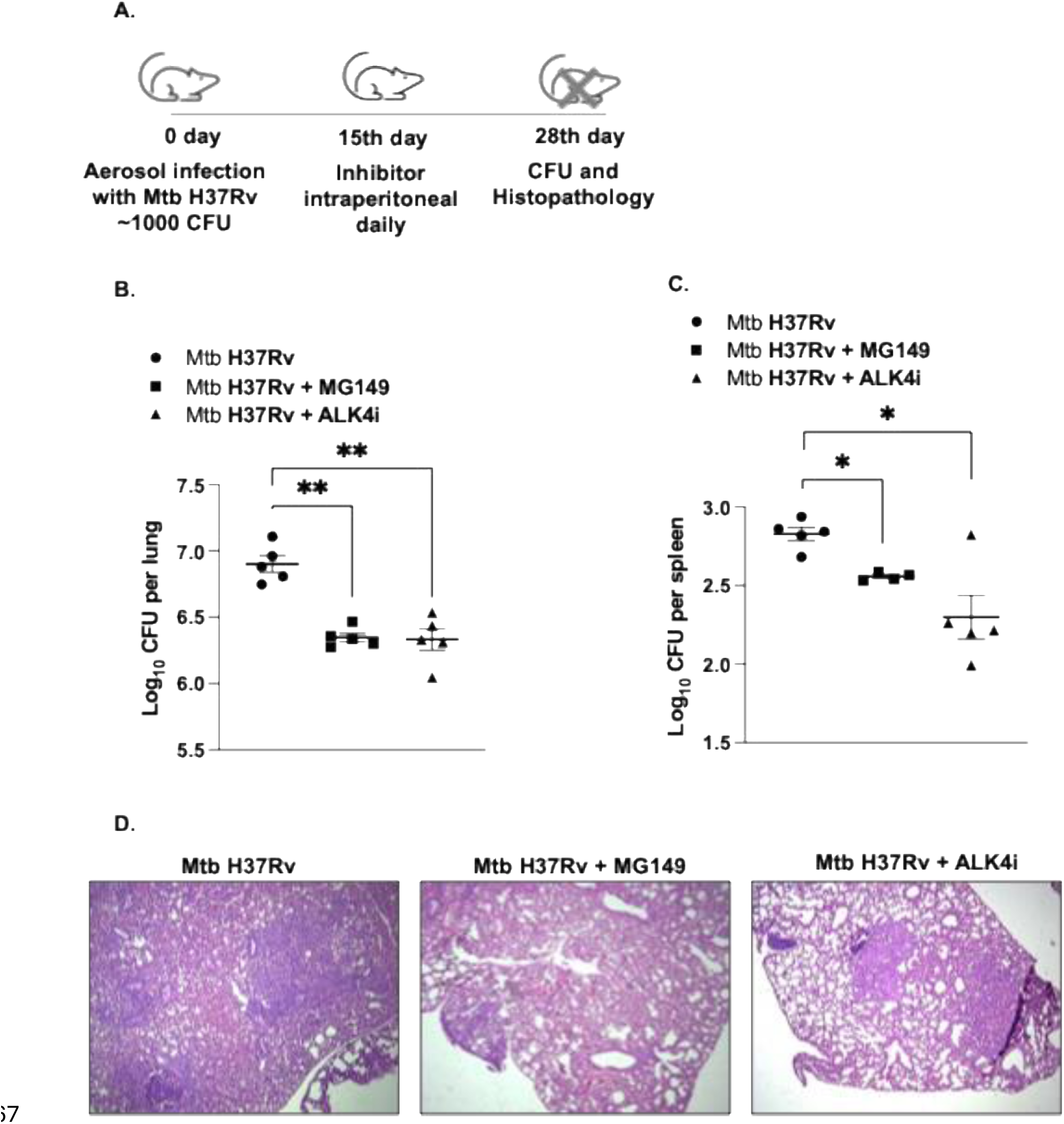
Inhibition of activin signalling and KAT8 alleviates TB pathology. **(A)** BALB/c mice were aerosol infected with Mtb H37Rv [∼1000 CFU] and treated with ALK4 inhibitor and KAT8 inhibitor (Number of mice per group = 5). Mycobacterial burden in the **(B)** lungs and **(C)** spleen of infected and inhibitor-treated mice was enumerated by plating homogenates on 7H11 plates. **(D)** TB pathology (granulomatous lesions) in the lungs was analysed by H and E staining. *, p < 0.05; **, p < 0.005 (Student’s t-test; GraphPad Prism 10.0).

Thus, we first report the role for activin A-mediated expression of the acetyltransferase, KAT8, in facilitating mycobacterial survival and dissemination. Increased KAT8 levels during Mtb infection acetylates the transcription factor, NRF2 contributing to its nuclear retention, and aiding in Mtb-induced ferroptosis. Together, we propose inhibitors targeting activin signalling and KAT8 as host-directed therapeutics that could assist in mycobacterial clearance as they target a crucial node used by Mtb for its dissemination and survival.

## Discussion

Mtb continues to remain a pertinent threat to human health and the leading cause of death due to infectious agents. With its ability to co-opt distinct host cellular proteins and signalling pathways to facilitate immune evasion, Mtb can persist within its host and resist killing by anti-mycobacterial drugs. In this regard, recent efforts have aimed at identifying targets for host-directed therapy wherein key nodes aiding in mycobacterial pathogenesis are targeted to shorten therapy time and improve treatment prospects. In this study, we showed that activin signalling-dependent expression of KAT8 regulates the activity of the master transcriptional regulator, NRF2, and induces ferroptosis by regulating the expression of HO-1. We observed a significant reduction in Mtb dissemination and survival upon utilizing specific inhibitors targeting these crucial nodes during Mtb infection. In this light, we believe targeting these factors would provide clinically relevant adjuncts for TB treatment.

As an intracellular pathogen, Mtb survival depends on the viability of its host cell. However, previous reports have indicated the role of necrosis and ferroptosis as a mechanism of Mtb release required for subsequent dissemination (16, 19). While several antioxidant mechanisms, including GPX4 and BACH1, have been reported to regulate Mtb-induced ferroptosis, the role of iron overload have not been reported during tuberculosis. Our findings provide the first evidence supporting the role of activin A in regulating the expression of HO-1 leading to Mtb-induced ferroptosis. While our study does not delineate the specific role of HO-1 induced heme degradation and the subsequent increase in LIP, previous reports have identified large quantities of intracellular free heme due to massive haemorrhage during active TB that could serve as the source of iron required for the induction of ferroptosis. Further, HO-1, the rate-limiting enzyme in heme catabolism, has also been reported to contribute to an increased labile iron pool during Mtb infection and has been identified as a marker for disease severity during tuberculosis (26, 27). However, studies have also indicated towards a dual role for HO-1 in TB pathogenesis. Reports on HO-1 deficient mice have demonstrated a protective role for HO-1 during TB, wherein deficient mice were more susceptible to Mtb infection, had increased bacterial loads, and higher mortality rates(39). Additionally, a parallel study has also demonstrated the role of HO-1 in granuloma formation and Mtb dissemination using murine models of TB(40). While reports have hypothesized that the excessive free heme released during later stages of tuberculosis could overwhelm the cytoprotective effects of HO-1 and contribute to oxidative stress and excessive ROS production (34), studies on their the overall impact are imperative towards understanding their distinct role in TB. Additionally, Mtb infection is also known to modulate other cell death pathways including apoptosis and pyroptosis (15). Furthermore, activin-A has been implicated in regulating these modes of cell death in distinct models of study (41). Consequently, we cannot exclude the possible role of Mtb-induced activin-A in regulation of other cell death pathways in TB.

In addition, we showed that Mtb infection induces the activin-A dependent activation of SMAD2/3 signalling which further regulates levels of the acetyltransferase, KAT8. While we report that KAT8 acetylates NRF2 leading to its nuclear localisation, its role in the regulation of other transcription factors and cellular processes during Mtb infection remains unknown. Parallel studies have also identified the role of KAT8 in inflammatory cytokine production and fibrosis which are implicated to contribute to disease pathology in TB (42, 43). Additionally, KAT8 has also been widely reported to regulate gene expression by transferring acetyl groups onto lysine residues on histones, loosening the chromatin structure and promoting gene transcription. Further, the BET protein, BRD4 has also been identified as a co-factor of KAT8 in regulating transcriptional activation thereby regulating autophagy and stress responses (44, 45). BRD4 has also been previously established to aid in the pathogenesis of Mtb (46) and the increased levels of KAT8 could be an additional mechanism of Mtb-mediated regulation of BRD4 activity. Thus, additional consequences of Mtb-mediated upregulation of KAT8, other than its role in regulating the nucleocytoplasmic localisation of NRF2, would be promising avenues of future research. Taken together, we report the novel role of Mtb-induced activin signalling and its downstream effector SMAD2/3 in regulation of KAT8. Additionally, we uncover the KAT8-mediated regulation of NRF2 acetylation and HO-1 expression in Mtb infected macrophages thereby inducing ferroptosis during Mtb infection. Future experiments with gene-specific knockout animals and detailed characterisation of the consequence of KAT8 expression would help in advancing our understanding of mycobacterial pathogenesis.

## Materials and Methods

### Mice and Cells

BALB/c mice, four-to six-week-old, male and female, were used during this study. Mice were purchased from The Jackson Laboratory and maintained in the Central Animal facility (CAF) at the Indian Institute of Science (IISc) under a 12h light and dark cycle. All in vitro experiments were performed in peritoneal macrophages isolated from these mice. Briefly, mice were injected intraperitoneally with 8% Brewer’s thioglycolate medium (HiMedia, M019) and were maintained in CAF for four days, after which they were sacrificed. Peritoneal exudates were harvested in ice-cold PBS, and adherent cells were utilized as peritoneal macrophages for experiments. Cells were cultured in Dulbecco’s Modified Eagle Medium (DMEM; Gibco, Thermo Fisher Scientific,12100061) supplemented with 10% heat-inactivated fetal bovine serum (FBS; Gibco, Thermo Fisher Scientific,10270106) and maintained at 37°C in 5% CO2 incubator.

### Ethics statement

Approval from the Institutional Ethics Committee for animal experimentation, Indian Institute of Science, Bengaluru, was obtained before the mice experiments were conducted. The animal care and use protocol adhered to was approved by national guidelines of the Committee for the Purpose of Control and Supervision of Experiments on Animals (CPCSEA), Government of India. Experiments with virulent mycobacteria (Mtb H37Rv) were approved by the Institutional Biosafety Committee.

### Bacteria

Virulent strain of Mycobacterium tuberculosis (Mtb H37Rv) was a kind research gift from Prof. Amit Singh, Centre for Infectious Disease Research, Microbiology and Cell Biology, IISc, Bengaluru. Briefly, a single colony of Mtb H37Rv was inoculated in Middlebrook 7H9 (Difco, USA, 271310) culture medium supplemented with 10% OADC (Oleic acid, Albumin, Dextrose, Catalase). At the mid-log phase of bacterial culture, single-cell suspension of mycobacteria was obtained by passing the culture through 23-, 26- and 30-gauge needles, ten times each. These suspensions were used to infect peritoneal macrophages at the multiplicity of infection 10. All the experiments involved with Mtb H37Rv were carried out in a Biosafety level 3 (BSL-3) facility at the Centre for Infectious Disease Research (CIDR), IISc, Bengaluru.

### *In vivo* infection of mouse

Madison chamber aerosol generation instrument calibrated to 1000 CFU/animal and was used to infect BALB/c mice with mid-log phase Mtb H37Rv. Aerosolized animals were maintained in a securely commissioned BSL3 facility. At 15 days of infection, mice were administered SB431542 (Cayman Chemical Company, 13031) (10mg/kg of mice) or MG149 (Cayman Chemical Company, Cay22135-5) (4mg/kg of mice) daily via intraperitoneal injections. Mice were sacrificed on the 28th day after infection. Specific lobes from the lungs of mice were isolated for histopathology studies and the extraction of RNA and protein.

### Treatment with pharmacological inhibitors

Mouse peritoneal macrophages were treated with the following reagents one hour prior to infection with mtb H37Rv. SB431542 (Cayman chemical company, 13031; 10 μM); ML385 (MedChemExpress, HY-100523; 5 μM); recombinant Follistatin (R & D system, 769-FS; 50ng/ml of medium).

### Transient Transfection

For siRNA transfection, mouse peritoneal macrophages were treated with 100nM of non-targeting siRNA or specific siRNA for Kat8 (Dharmacon, USA), Alk4 and Hmox-1 (Eurogentec) with the help of polyethyleneimine (PEI; Sigma-Aldrich, 764,604) or Lipofectamine 3000 (Thermo fisher scientific, Invitrogen, L3000015) for 6 h; followed by 24 h recovery. Transfected cells were then subjected to the infection for indicated time points and processed for analysis.

### Immunoblotting

For immunoblotting, cells were washed with PBS after treatment or infection. Whole-cell lysate was prepared by lysing the cells in RIPA buffer [50 mM Tris-HCl (pH 7.4), 1% NP-40, 0.25% sodium deoxycholate, 150 mM NaCl, 1 mM EDTA, 1 μg/ml each of, pepstatin, leupeptin, aprotinin, 1 mM NaF, 1 mM Na3VO4] on ice for 30 min. The total protein from cell lysates was estimated by the Bradford reagent. Subsequently, an equal amount of protein was resolved on 12% or 10% SDS-PAGE and transferred onto PVDF membranes (Millipore, IPVH00010) by semi-dry immunoblotting method (Bio-Rad). Blots were blocked using 5% non-fat dry milk powder in TBST [20 mM Tris-HCl (pH 7.4), 137 mM NaCl, and 0.1% Tween 20] for 60 min at room temperature. After blocking, blots were washed using TBST and probed with primary antibody diluted with 5% BSA in TBST overnight at 4°C. After washing with TBST, blots were probed with an anti-rabbit secondary antibody conjugated with HRP for 4h at 4°C. The immunoblots were developed with a western ECL detection system (Clarity, Bio-Rad, 1705061) per the manufacturer’s instructions. For developing more than one protein at a particular molecular weight range, the blots were stripped off the first antibody at 60 °C for 5 min using stripping buffer (62.5 mM Tris-HCl, with 2 % SDS, 100 mM 2-Mercaptoethanol), washed with 1X TBST, blocked (as mentioned above); followed by probing with the subsequent antibody following the described procedure. ACTB was used as a loading control.

### RNA isolation and quantitative real-time PCR (qRT-PCR)

Cells after infection or treatment were harvested using TRI-reagent (Sigma-Aldrich, T9424), and chloroform was added for phase separation. Total RNA was precipitated from the aqueous layer. RNA was used for cDNA synthesis using a first-strand cDNA synthesis kit (Promega, M3682). Quantitative real-time PCR was performed with SYBR green PCR mix (Takara, RR420A). *Gapdh* has been used as an internal control. Primer pairs used for expression analysis are provided in Supplementary Table 1.

### Bacterial enumeration

Cells were infected with Mtb H37Rv at MOI 10 for 4h. After 4h of infection, macrophages were washed thoroughly with PBS to eliminate the presence of the extracellular bacterium. Cells were then supplemented with a medium containing amikacin (HiMedia, CMS644-1G) at 0.2mg/ml for 2h to deplete any extracellular bacterium. Cells were washed with PBS and supplemented with DMEM medium at this time, taken as 0h. Duplicates were maintained with respective inhibitors in the antibiotic-free medium. The extracellular mycobacterial burden was enumerated by platting the medium with appropriate dilution on Middlebrook 7H11 (Difco, USA, 212203) supplemented with 10% OADC. Total colony-forming units (CFUs) were counted after 21 days.

### Chromatin Immunoprecipitation Assay

The chromatin immunoprecipitation (ChIP) assays were carried out using the protocol provided by Sigma-Aldrich and Abcam with certain modifications. In brief, infected or treated samples were washed with ice-cold phosphate-buffered saline, and samples were fixed with 3.6% paraformaldehyde solution for 15 mins at room temperature, followed by the inactivation of paraformaldehyde using 125mM glycine. Nuclei were lysed in 0.1% Sodium dodecyl sulfate (SDS) lysis buffer (50 mM Tris-HCl [pH 8.0]; 200 mM sodium chloride [NaCl]; 10 mM HEPES [pH 6.5]; 0.1% SDS; 10 mM ethylenediaminetetraacetic acid [EDTA]; 0.5 mM ethylene glycol-bis(β-aminoethyl ether)-N, N, N′, N′-tetraacetic acid; 1 mM phenylmethylsulfonyl fluoride; 1 μg/mL each of aprotinin, leupeptin, and pepstatin; 1 mM sodium orthovanadate; and 1 mM Sodium fluoride). The chromatin shearing Bioruptor Plus device (Diagenode, Belgium) was used at high power for 60 cycles with a 30-second pulse on and 45 seconds off. Chromatin extract containing DNA fragments with an average size of 500 bp was immunoprecipitated with indicated antibodies (Supplementary Table 2) or incubated with (Pierce, Thermo Fisher Scientific, 88803) Protein A/G Magnetic Beads. Immunoprecipitated complexes were washed with Wash Buffer A, Wash Buffer B, and TE (Wash Buffer A: 50 mM Tris-HCl [pH 8.0], 500 mM NaCl, 1 mM EDTA, 1% Triton X-100, 0.1% sodium deoxycholate, 0.1% SDS, and protease/phosphatase inhibitors; Wash Buffer B: 50 mM Tris-HCl [pH 8.0], 1 mM EDTA, 250 mM lithium chloride, 0.5% NP-40, 0.5% sodium deoxycholate, and protease/phosphatase inhibitors; TE: 10 mM Tris-HCl [pH 8.0], 1 mM EDTA) and followed by elution using elution buffer (1% SDS, 0.1 M sodium bicarbonate). Eluted samples were treated with RNase A and Proteinase K, followed by DNA purification and precipitation using the phenol-chloroform-ethanol method. Purified DNA was analysed by qRT-PCR. All values in the test samples were normalized to amplification of the specific gene in Input and IgG pull-down and represented as fold change in modification or enrichment. All ChIP experiments were repeated at least 3 times. The primer pairs used for the expression analysis are provided in Supplementary Table 3.

### Immunoprecipitation Assay

Immunoprecipitation assays were carried out following a modified version of the protocol provided by Millipore, USA. In brief, cells were washed with ice-cold PBS after treatment or infection, followed by lysis using RIPA buffer. Samples were pre-cleared using the BSA-blocked Protein A beads (Merck Millipore, 16-125) for 30 mins at 4℃. Cell lysates were then quantified, and an equal amount of protein was used for the pull-down from each treatment with antibody of interest or IgG isotype O/N at 4℃ on rotation. Protein A beads were added to the sample for 4h at 4℃. The pellet containing beads bound immune complexes were then washed using RIPA buffer, and protein complexes were eluted by boiling the beads in Laemmli buffer for 10 min. The bead-free samples were resolved by SDS-PAGE and the target interacting partners were identified by immunoblotting. Clean-Blot™ IP Detection Reagent (Pierce, Thermo Fisher Scientific, 21230) is used as a secondary antibody for immunoblotting after immunoprecipitation was obtained from Thermo Scientific.

### Immunofluorescence and lipid staining

Cells were fixed with 3.7 % formaldehyde. Fixed samples were blocked with 2 % BSA containing 0.02 % saponin (for intracellular staining) for 1 h at room temperature. After blocking, the cells were stained with primary antibody (diluted 1:250 in respective blocking solution) at 4 °C. Following overnight incubation, samples were washed thoroughly with 1X PBS and incubated with fluorescent dye-conjugated secondary antibody DyLight 488- (Thermo Fisher Scientific, 35502) for 2 h at room temperature. Samples were washed and stained with DAPI (Sigma-Aldrich, D9542) for 10 min at room temperature to mark the nuclei. Formaldehyde-fixed cells were stained with BODIPY 493/503 (Invitrogen, D3922) and BODIPY 665/676 (Invitrogen, B3932) for lipid peroxidation staining. Samples were then stained to mark the nuclei using DAPI for 10 mins at room temperature. After thorough washing, samples were mounted on glycerol for visualization by confocal microscopy [Zeiss LSM 710 Meta confocal laser scanning microscope (Carl Zeiss AG, Germany) using a plan Apochromat 63X/1.4 Oil DIC objective (Carl Zeiss AG, Germany)] and analysis using ZEN 2009 software.

### Detection of intracellular Labile Iron Pool

Peritoneal macrophages were seeded in a 96-well plate. Cells after treatment or infection were washed using 1X PBS. Cells were then incubated with FerroOrange dye (Merk Millipore, SCT-210) for 30 mins in a 37℃-incubator equilibrated with 95% air and 5% CO2, and the cells were washed with PBS (200 μl) three times. Cells were analyzed using plate reader SpectraMax M3 at Ex/Em = 543 nm/ 580 nm.

### Cell death measurement

Cellular necrosis was measured by LDH release. Briefly, Mouse peritoneal macrophages were seeded in 96well plate cells and were pre-treated with indicated inhibitors before mtb H37Rv infection for the timepoint mentioned in the results, and LDH release was measured by CyQUANT LDH Cytotoxicity Assay (Invitrogen, C20300) according to the manufacturer’s instructions, and absorbance was measured using plate reader SpectraMax M3.

### Statistical analysis

Levels of significance for comparison between samples were determined by the student’s t-test and one-way ANOVA followed by Tukey’s multiple-comparisons. The data in the graphs are expressed as the mean ± S.E.M for the values from at least 3 or more independent experiments and P values < 0.05 were defined as significant. GraphPad Prism software (version 10.0, GraphPad Software, USA) was used for all the statistical analyses.

## Supporting information

Supplemental Informantion

## Acknowledgments

We thank the Central Animal Facility, Indian Institute of Science (IISc), for maintaining and providing mice for experimentation. We acknowledge the help of the Biosafety Level 3 facility and staff at Centre of Infectious Diseases (CIDR), IISc, for helping us in our *in vitro* and *in vivo* experiments with Mtb H37Rv. We are also grateful to the Department of Microbiology and Cell Biology confocal facility. We are grateful to Prof. K. N. Balaji’s research group, majorly Awantika Shah for their valuable inputs in improving the manuscript

## Footnotes

This work was supported by the Department of Biotechnology, Government of India (DBT number BT/PR47843/MED/29/1631/2023, DT. .25.09.2024; BT/PR41341/MED/29/1535/2020 DT. 13.08.2021; BT/PR27352/BRB/10/1639/2017, DT.30/8/2018; and BT/PR13522/COE/34/27/2015, DT.22/8/2017 to K. N. B.) and the Department of Science and Technology, Government of India (DST) (EMR/2014/000875, DT.4/12/15 to K. N. B.), New Delhi, India. K. N. B. thanks the Science and Engineering Research Board (SERB), DST, for the award of the J. C. Bose National Fellowship (number SB/S2/JCB-025/2016 dt 25.7.15), second term J. C. Bose National Fellowship (JBR/2021/000011), and core research grant (CRG/2019/002062). The authors thank DST-Fund for Improvement of S&T Infrastructure, University Grants Commission (UGC) Centre for Advanced Study, and DBT-IISc Partnership Program (phase II at IISc, BT/PR27952/INF/22/212/2018) for funding and infrastructure support. K. N. B. also acknowledges the funds received as a part of infrastructure support to IISc as a part of the Institute of Eminence (IOE) scheme of the Government of India. Fellowships were received from UGC and IISc (B. A. S.), CSIR and The Prime Minister’s Research Fellows Scheme (to S. S.). The funders had no role in study design, data collection and analysis, decision to publish or preparation of the manuscript.

## Author Contributions

B.A.S and K.N.B. conceived and designed the study. B.A.S, S.S, and performed the experiments and analysed the data. R.S.R., B.A.S and S.S performed the animal experiments. B.A.S, S.S and K.N.B. wrote the manuscript.

