## Supplemental Informantion for "Activin A mediated KAT8 expression induces ferroptosis during *Mycobacterium tuberculosis* infection"

**Supplemental Material**

**Table S1: List of primers used in the study for gene expression analysis**

| **Genes** | **Forward primer (5'-3')** | **Reverse primer (5'-3')** |
|---|---|---|
| *Gapdh* | GAGCCAAACGGGTCATCATCT | GAGGGGCCATCCACAGTCTT |
| *Inhba* | TGAGAGGATTTCTGTTGGCAAG | TGACATCGGGTCTCTTCTTCA |
| *Inhbb* | CTTCGTCTCTAATGAAGGCAACC | CTCCACCACATTCCACCTGTC |
| *Kat8* | ACGAGGCGATCACCAAAGTG | AAGCGGTAGCTCTTCTCGAAC |
| *Hmox1* | CACGCATATACCCGCTACCT | CCAGAGTGTTCATTCGAGA |

**Table S2: List of antibodies used in the study**

| **Antibody** | **Company** | **Catalogue No** |
|---|---|---|
| ALK4 | Abcam | ab109300 |
| pSMAD2 | Invitrogen | 442449 |
| pSMAD3 | Invitrogen | 443469 |
| KAT8 | Abcam | Ab200660 |
| NRF2 | CST | 12721 |
| HO-1 | Abcam | Ab68477 |
| HRP-tagged anti-β-ACTIN | Sigma | A3854 |
| HRP conjugated anti-rabbit IgG | Jackson ImmunoResearch (USA) | 111–035-045 |

**Table S3 : List of primers used for ChIP analysis**

| **Genes** | **Forward primer (5’-3’)** | **Reverse primer (5’-3’)** |
|---|---|---|
| NRF2 binding site | | |
| *Hmox1* | GCTGGAATGCTGAGTTGTGA | TGAGGGAACAGAGGGTGACT |
| pSMAD3 binding site | | |
| *Kat8* site 1 | AGTTTACCTTGAATGCTGCT | GACACTCCTCCAAGTACAGA |
| *Kat8* site 2 | GCCAGGGTTATACAGAGGA | GTCAGAGTAGGGCTGGATAA |

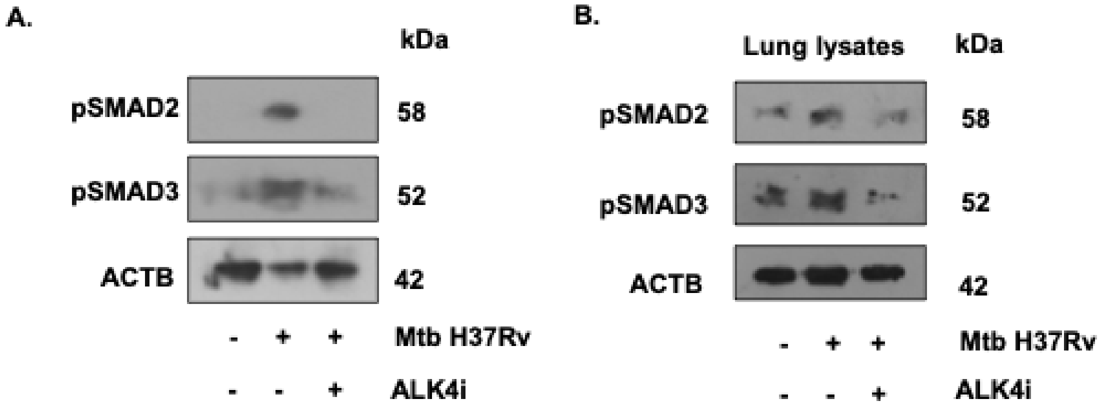


**Figure S1** (A) Mouse macrophages were pre-treated with ALK4 inhibitor (SB431542) and infected with Mtb H37Rv for 60 mins. Whole cell lysates were assessed for phosphorylation of SMAD2/3 using immunoblotting. (B) Lung lysates from BALB/c mice aerosol-infected (~1000 colony-forming units [CFU]) with Mtb H37Rv and administered with ALK4 inhibitor (SB-431542) from 15 d p.i were assessed for the for phosphorylation of SMAD2/3 at 28 d p.i. All immunoblotting data is representative of three independent experiments. ACTB was utilized as a loading control.

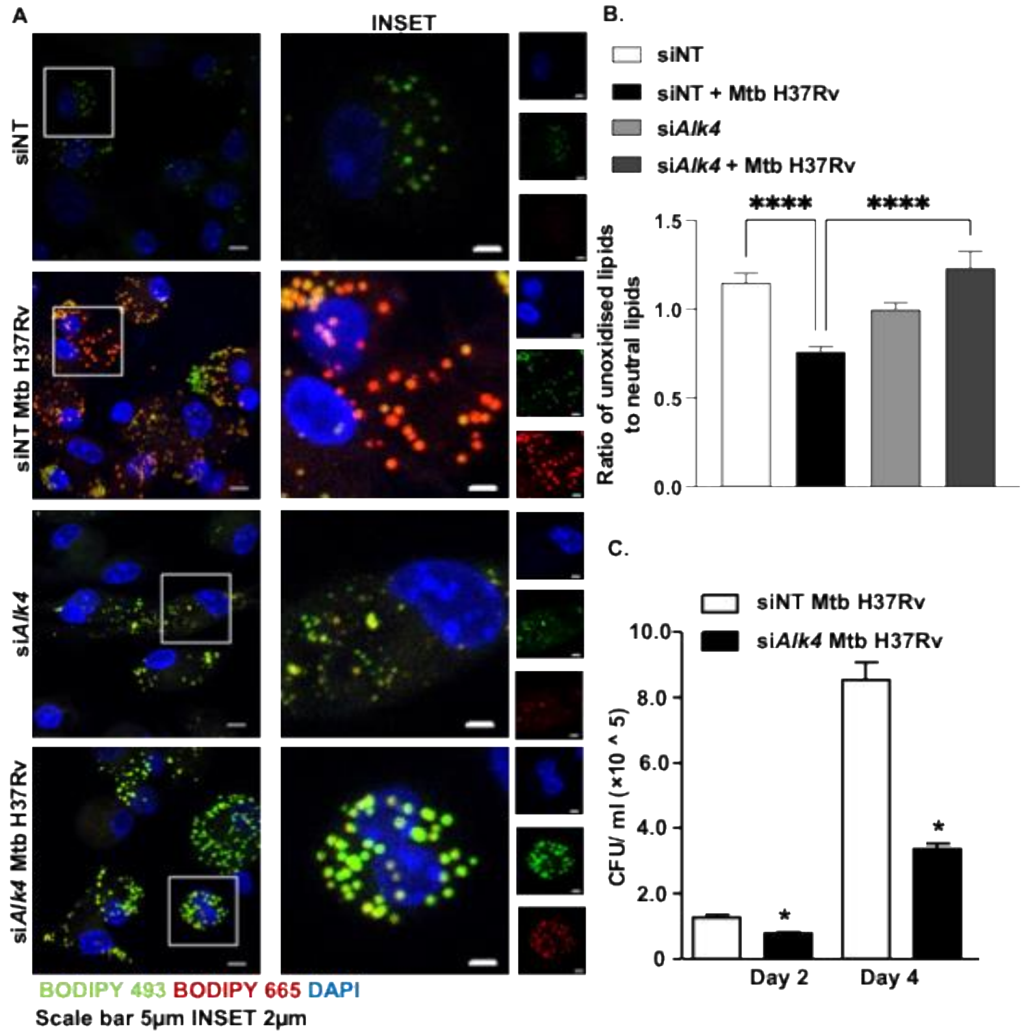


**Figure S2: (A, B)** Mouse macrophages were transfected with non-targeting or *Alk4* small interfering RNAs. Transfected cells were infected 24 h with Mtb H37Rv, and lipids were stained with BODIPY 493/503 and BODIPY 665/676 to assess for lipid peroxidation and observed by confocal microscopy; representative images (A) and its respective quantification (B). **(C)** Mouse macrophages were transfected with non-targeting or *Alk4* small interfering RNAs. The release of live mycobacteria from necrotic cells was examined by CFU quantification in transfected cells by plating culture supernatants 48 h and 96 h post-infection. *, $p < 0.05$; ****, $p < 0.0001$ (Student's t-test in C; One-way ANOVA in B; GraphPad Prism 10.0).

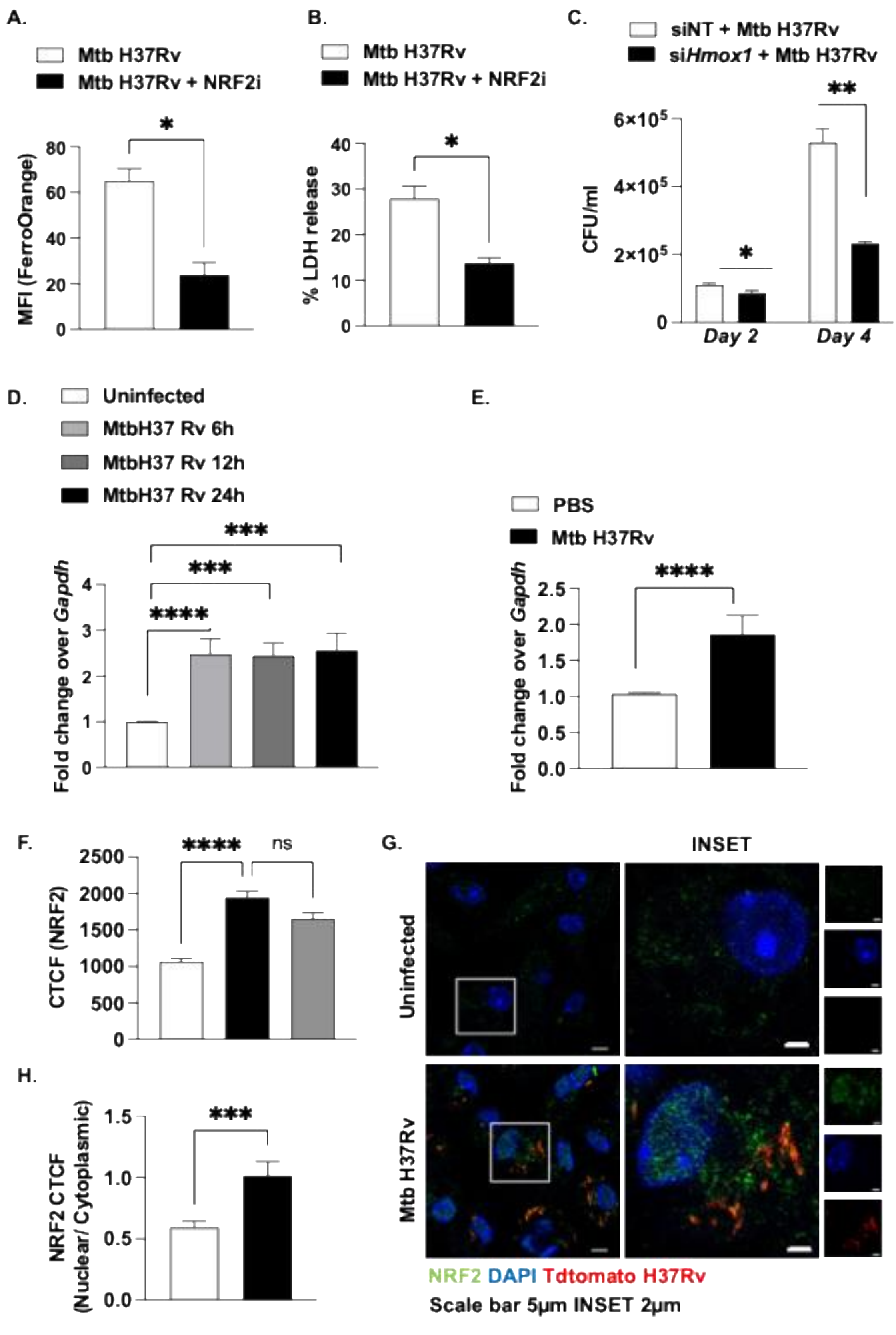


**Figure S3: (A)** Mouse macrophages were pre-treated with NRF2 inhibitor (ML385) followed by 24 h infection with Mtb H37Rv. The labile iron pool was quantified via staining with FerroOrange. **(B)** Necrotic cell death was measured by LDH release assay in mouse macrophages treated with NRF2 inhibitor (ML385) followed by Mtb H37Rv infection for 96 h. **(C)** Mouse macrophages were transfected with non-targeting or *Hmox1* small interfering RNAs. The release of live mycobacteria from necrotic cells was examined by CFU quantification in transfected cells by plating culture supernatants 48 h and 96 h post-infection **(D)** Mouse macrophages were infected with Mtb H37Rv for indicated time points and assessed for transcript levels of HO-1 using qRT-PCR. **(E)** BALB/c mice were aerosol-infected (∼1000 colony-forming units [CFU]) with Mtb H37Rv for 28 d, and transcript levels of HO-1 were assessed in RNA isolated from the lungs of uninfected and infected mice using qRT-PCR. **(F)** Mouse macrophages were pre-treated with ALK4 inhibitor (SB-431542), infected with Mtb H37Rv for 24 h and assessed for levels of NRF2 by confocal microscopy **(G, H)** Mouse macrophages were infected with Mtb H37Rv for 24 h and assessed for nuclear levels of NRF2 by confocal microscopy representative image **(G)** its respective quantification **(H)**. All confocal microscopy data are representative of three independent experiments. *, $p< 0.05$; **, $p< 0.005$; ***, $p < 0.0005$; ****, $p < 0.0001$ (Student's t-test in A, B, C, D, E, H; One-way ANOVA in F; GraphPad Prism 10.0).

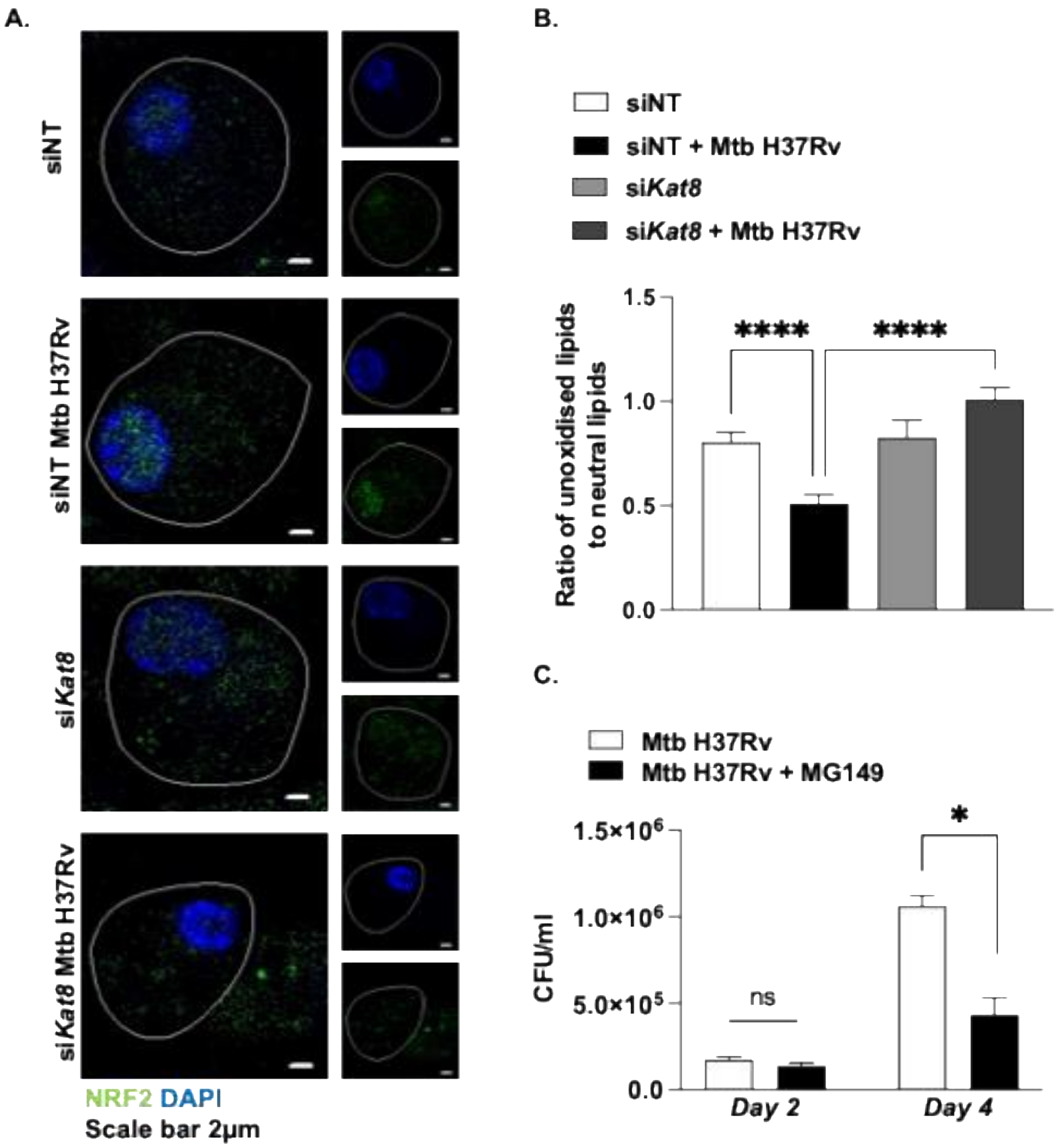


Figure S4: **(A)** Mouse macrophages were transfected with non-targeting or *Kat8* small interfering RNAs. Transfected cells were infected 24 h with Mtb H37Rv, and assessed for levels of NRF2 using confocal microscopy **(B)** Mouse macrophages were transfected with non-targeting or *Alk4* small interfering RNAs. Transfected cells were infected 24 h with Mtb H37Rv, and lipids were stained with BODIPY 493/503 and BODIPY 665/676 and lipid peroxidation was quantified. **(C)** The release of live mycobacteria from necrotic cells was examined by CFU quantification in mouse macrophage pre-treated with MG149 (KAT8 inhibitor) by plating culture supernatants 48 h and 96 h post-infection.
